# PATJ regulates cell stress responses and vascular remodeling post-stroke

**DOI:** 10.1101/2024.07.17.603777

**Authors:** Mengqi Zhang, Wei Jiang, Kajsa Arkelius, Raymond A. Swanson, Dengke K. Ma, Neel S. Singhal

## Abstract

PALS1-associated tight junction (PATJ) protein is linked to metabolic disease and stroke in human genetic studies. Despite the recognized role of PATJ in cell polarization, its specific functions in metabolic disease and ischemic stroke recovery remain largely unexplored. Using a mouse model of stroke, we found post-ischemic stroke duration-dependent increase of PATJ abundance in endothelial cells. PATJ knock-out (KO) HEK293 cells generated by CRISPR-Cas9 suggest roles for PATJ in cell proliferation, migration, mitochondrial stress response, and interactions with the Yes-associated protein (YAP)-1 signaling pathway. Notably, *PATJ* deletion altered YAP1 nuclear translocation. *PATJ* KO cells demonstrated extensive transcriptional reprograming based on RNA sequencing analysis. Crucially, we identified dysregulation in genes central to vascular development, stress response, and metabolism, including *RUNX1*, *HEY1*, *NUPR1*, and *HK2*. These insights offer a new understanding of PATJ’s complex regulatory functions within cellular and vascular physiology and help lay the groundwork for therapeutic strategies targeting endothelial PATJ-mediated pathways for stroke rehabilitation and neurovascular repair.

## 1. Introduction

Stroke is a principal cause of mortality and prolonged disability in the United States. While thrombolysis with tissue-type plasminogen activator (tPA) or endovascular thrombectomy has improved outcomes following AIS [1], strategies to improve outcomes for stroke survivors are still notably lacking [2]. Angiogenesis, the process of new blood vessel formation from existing ones is tightly regulated under normal physiological conditions in adults [3], however, following AIS, angiogenesis is a critical adaptation restoring cerebral blood flow and supporting repair processes [4–6]. Interestingly, an increased density of microvessels in brain regions impacted by AIS correlates with better prognosis in patients [7], thus promoting angiogenesis following AIS is a promising method for enhancing neurorecovery from AIS.

The evolutionarily conserved Crumbs protein homologs (CRB) are key components in scaffolding and cellular orientation. Central to this orchestration is the PALS1-associated tight junction (PATJ) protein, a critical component in the CRB complex. PATJ serves as a connector between the CRB– PALS1 complex and tight junction proteins [8, 9]. This role is critical in maintaining cell polarization and ensuring the stability of apical junctions [10, 11]. However, recent advancements have begun to unveil a broader spectrum of PATJ’s functionality including roles in signal transduction, cellular dynamics, and tumorigenesis [12]. A recent genome-wide association study (GWAS) on acute ischemic stroke (AIS) outcomes revealed that AG or GG single nucleotide polymorphisms (SNPs) of the PATJ gene are strongly associated with poor 90-day modified Rankin Scale (mRS) outcomes post-AIS [13]. Interestingly, SNPs in PATJ have also been implicated in cardiometabolic health and sleep disturbances [14, 15]. These genome-scale findings provide critical genomic links between PATJ and cardiovascular pathology, suggesting that variations in PATJ not only impact cellular polarity but may also influence cardiovascular risk and AIS recovery outcomes.

*PATJ* knockdown in human microvascular endothelial cells was found to promote endothelial to mesenchymal transition and pro-angiogenic programs [16]. Prior work indicates an important role for PATJ in directing embryonic capillary migration in cooperation with angiomotin (AMOT) and *Syx*, a RhoA GTPase exchange factor (RhoGEF) protein also known as Pleckstrin homology domain containing, family G member (*PLEKHG*)-5 [17]. The involvement of endothelial cells in vascular remodeling and angiogenesis is well-documented, yet the specific contributions of PATJ during post-AIS angiogenesis remains incompletely understood. Our study builds upon this recent expanded understanding of PATJ, and links PATJ with post-AIS angiogenesis, cellular migration, proliferation, and metabolism. We also found regulatory interplay between PATJ and key signaling pathways, notably YAP1 within the Hippo pathway. Taken together, these findings highlight PATJ’s role in neurorepair-related pathways and suggest its potential as a novel therapeutic target.

## 2. Methods

### 2.1 CRISPR-Cas9 RNP complex preparation and cell transfection

For the targeted editing of *PATJ* exon 3, sgRNA sequences sgRNA1 (CUUGACCCAAGAUAAACUGC) and sgRNA2 (GUGAGUAUCUGGUUGAAGAG) were synthesized with high specificity and minimal off-target effects (Synthego). The sgRNAs were then complexed with Cas9 protein (IDT) to form ribonucleoprotein (RNP) complexes. The complexes were prepared by diluting sgRNA and Cas9 to 3 μM in Opti-MEM™ I Reduced Serum Medium, adhering to a sgRNA:Cas9 molar ratio of 1.3:1, followed by a brief incubation.

HEK293 cells, cultured in Dulbecco’s Modified Eagle Medium (DMEM) supplemented with 10% Fetal Bovine Serum and antibiotics, were grown to 30-70% confluence in 24-well plates to ensure optimal cell density for transfection. The transfection utilized Lipofectamine™ CRISPRMAX™ (Invitrogen) for RNP complex delivery, employing a reverse transfection technique where the RNP solution is pre-mixed with the cell suspension before plating. Following transfection, the cells were incubated at 37°C and 5% CO2, with medium changes to maintain cell health and promote recovery.

### 2.2 Isolation and validation of CRISPR-edited clones

Following transfection, single-cell derived clones were derived from singly-plated cells, ensuring genetic uniformity. Over 2-8 weeks, these colonies were observed under standard conditions and expanded based on visual confirmation via microscopy. Successfully expanded colonies were then transferred to 24-well plates for further growth. Sanger sequencing analysis post-PCR amplification from genomic DNA of individual clones and Western blot was performed to validate targeted knockout of *PATJ* exon 3.

### 2.3 Migration assay

Confluent HEK293 cell monolayers, both *PATJ* knockout (KO) and wild-type (WT), were subjected to a scratch assay. A sterile pipette tip was used to create a straight ‘wound’ across the monolayer. Cell proliferation was inhibited using Mitomycin C (10µg/mL). The migration of cells into the wound area was imaged at specified intervals to assess closure rate. Quantitative analysis was conducted by measuring the gap width at 0, 24, and 48 hours post-scratch. Images were captured under an Zeiss AXIO Observer.A1 Inverted Fluorescence Microscope. Cellular migration was determined by ImageJ software. Relative migration rate was calculated as (Areawound at 0 h – Areawound at 24 h)/Areawound at 0 h.

### 2.4 Rotenone-induced cell growth assessment in PATJ-modified HEK293 cells

HEK293 cells, both with and without *PATJ* knockout, were cultured in 24-well plates at a seeding density of 10^5^ cells per well. Four experimental groups were established: PATJ KO and WT cells, each subjected to treatment with either 200 nM rotenone or the corresponding vehicle as a control. The rotenone concentration was selected to inhibit mitochondrial complex I, thereby simulating metabolic stress analogous to ischemic conditions. Cell proliferation was assayed by automated cytometry quantification of cell number at specific intervals post-treatment, directly within the culture wells (Nanoentek, Waltham, MA, USA).

### 2.5 Seahorse XF metabolic flux analysis

Mitochondrial respiration in HEK293 cells, with and without *PATJ* modification, was evaluated using the Seahorse XF^e^96 Extracellular Flux Analyzer (Seahorse Bioscience, North Billerica, MA, USA) with the Seahorse XF Cell Mito Stress Test Kit. Cells were plated at 25,000 cells per well in XF^e^96 microplates and incubated in standard culture medium. Following overnight incubation, cells were washed and equilibrated in unbuffered Seahorse XF Base Medium supplemented with glucose (4.5 g/L), L-glutamine (2 mM), and sodium pyruvate (1 mM) in a non-CO2 incubator at 37°C for 1 hour. Oxygen consumption rate (OCR) was recorded before and after sequential injections of mitochondrial inhibitors: oligomycin (1 μM), Trifluoromethoxy carbonylcyanide phenylhydrazone (FCCP; 1 μM), and rotenone/antimycin A (1 μM). Post-assay, the cells stained with Hoechst 33258 (0.1 μg/mL) and fluorescence intensity acquired at excitation/emission 355/460 nm to normalize the OCR data.

### 2.6 Assessment of stress responses in *C. elegans*

*C. elegans* strains were maintained on nematode growth medium (NGM) plates seeded with *Escherichia coli* OP50 at 20°C using standard procedures[18]. The *mpz-1(mib188[mpz-1::gfp])* II strain [19] was used in this study. For stress treatments, L4 stage worms were subjected to either hypoxia (0.1% O2) or normoxia for 72 hours, or heat stress at 20°C (control), 25°C, or 28°C for 72 hours[20]. For confocal imaging, treated MPZ-1::GFP worms were randomly selected, immobilized with 10 mM sodium azide in M9 buffer, and aligned on 2% agarose pads on glass slides. Images were acquired using a Leica TCS SPE confocal microscope with a 63× objective lens, maintaining consistent settings across all samples. MPZ-1::GFP abundance was quantified using ImageJ.

### 2.7 Western Blot

Protein lysates were extracted from HEK293 cells, both *PATJ* KO and WT, using cell lysis buffer (Cell Signaling Technology, Danvers, MA) supplemented with Protease Inhibitor Cocktail (Cell Signaling Technology, Danvers, MA). Equal amounts of protein were separated on 4-15% precast gels (Bio-Rad) and then transferred onto polyvinylidene difluoride (PVDF, Bio-Rad) membranes. Membranes were blocked for 1 hour at room temperature using Tris-buffered saline with 0.1% Tween-20 (TBS-T) supplemented with 5% non-fat dried milk. Membranes were then incubated overnight at 4°C with primary antibodies diluted in TBS-T containing 5% bovine serum albumin (BSA). Primary antibodies used were PATJ (1:1000, Proteintech) and YAP1(1:2000, Proteintech). Subsequently, membranes were probed with appropriate HRP-conjugated secondary antibodies for 1 hour at room temperature. Protein bands were visualized using an enhanced chemiluminescence (ECL) detection system (Azure Biosystems). Densitometric analysis was conducted to quantify protein expression levels, with α-tubulin employed as a loading control.

### 2.8 Cellular immunofluorescence

HEK293 cells, both *PATJ* KO and WT, were cultured on coverslips until optimal confluence was reached. Cells were fixed in 4% paraformaldehyde, then permeabilized with 0.1% Triton X-100 in PBS for 15 minutes. Non-specific binding was minimized by blocking with goat serum. Overnight incubation at 4°C with primary antibodies targeting YAP1 (1:200, Proteintech) was followed by application of goat anti-rabbit Alexa Fluor 488 secondary antibody (1:1000, Invitrogen). The cytoskeleton was stained with CoraLite 594-Phalloidin (red, Proteintech), and nuclei were stained with DAPI. Coverslips were mounted for examination by Leica confocal microscopy, focusing on YAP localization and expression. YAP localization was categorized as predominantly nuclear, cytoplasmic, or distributed across both compartments. Analysis included over 200 cells from each group in three separate experiments.

### 2.9 Mouse model of transient focal cerebral ischemia

Transient focal cerebral ischemia was induced in C57BL6 mice aged 8-10 weeks (weighing 23-26 g) using the Middle Cerebral Artery Occlusion (MCAO) technique. Mice were anesthetized with isoflurane, delivered in a mixture of oxygen and nitrous oxide. A midline neck incision was made to expose the common carotid artery, followed by electrocoagulation of its branches. A nylon filament was carefully inserted through the external carotid artery to the internal carotid artery, and advanced towards the middle cerebral artery (MCA) to achieve a reduction in regional cerebral blood flow (CBF) to less than 25% of the baseline values. Monitoring of the regional cerebral blood flow was conducted through Laser Doppler Flowmetry (Moor Instruments Ltd, UK). The mice’s body temperature was maintained at 37.0 ± 0.5°C using a regulated heating pad, monitored with a rectal thermometer throughout the procedures. The occlusion was sustained for 60 minutes, after which reperfusion was initiated. Mice were then allowed a recovery period ranging from 1 to 28 days. For the control group, sham operations were performed where all steps were identical except for the artery occlusion. Postoperative care included the administration of buprenorphine for pain relief. Mice failing to meet the specified CBF criteria or those that did not survive the procedure (n = 5) were excluded from subsequent analyses.

### 2.10 Immunohistochemistry

After transient Middle Cerebral Artery Occlusion (tMCAO), mice were anesthetized deeply and perfused transcardially, first with 0.9% NaCl, followed by 4% paraformaldehyde in PBS. Brains were then extracted, cryoprotected in 30% sucrose in PBS for 48 hours, and sectioned coronally on a cryostat. Sections were preserved at -20°C in a cryoprotectant solution until required for analysis or immunofluorescence staining. For staining, sections were washed in PBS, permeabilized with PBST (1% Triton X-100 in PBS) for 20 minutes and blocked with 5% normal goat serum for an hour. Following further washing with 0.3% PBST, they were incubated overnight at 4°C with rat anti-mouse CD31 (1:200, BD Pharmingen) and rabbit polyclonal PATJ antibody (1:100, LSBio). Alexa Fluor 488 conjugated goat anti-rat (1:1000, Invitrogen) or Alexa Fluor 594 goat anti-rabbit (1:1000, Invitrogen) secondary antibodies were applied respectively. For NeuN/PATJ staining, sections underwent incubation with NeuN (1:1000, LSBio) and PATJ antibodies, followed by the appropriate Alexa Fluor-conjugated secondary antibodies. Immunofluorescence images were captured using a Zeiss AXIO Observer.A1 Inverted Fluorescence Microscope. For YAP1/CD31/DAPI triple staining, sections were incubated overnight at 4°C with rat anti-mouse CD31 (1:200, BD Pharmingen) and rabbit polyclonal YAP1 antibody (1:200, Proteintech). Alexa Fluor 594 conjugated goat anti-rat (1:1000, Invitrogen) and Alexa Fluor 488 goat anti-rabbit (1:1000, Invitrogen) secondary antibodies were applied respectively. Sections were then mounted with VECTASHIELD Vibrance® Antifade Mounting Medium with DAPI (H-1800) for nuclear counterstaining. Images were acquired using a CSU-W1 SORA spinning disk confocal microscope.

### 2.11 RNA-seq analysis of HEK293 cell lines

Total RNA was extracted from HEK293 cells across three experimental groups: *PATJ* KO, WT, and *PATJ* KO with overexpression, utilizing the RNeasy Mini Kit (Qiagen). Prior to library preparation, the integrity and concentration of RNA were verified. Sequencing libraries were constructed following cDNA synthesis, adaptor ligation, and amplification. Library quality control followed by next-generation sequencing was performed on the DNBSeq platform by a commercial provider (BGI Americas, San Jose, CA). Reads were aligned to the human genome (Hgv38) for differential gene expression analysis using the limma R package (version 3.40.6; [21]), which utilizes generalized linear models. Volcano plots and heatmaps were generated with the ggplot2 (version 3.4.0;[22]) and pheatmap (version 1.0.12) R packages, respectively. For dimensionality reduction, the umap R package (version 0.2.7.0) was used, normalizing expression profiles (z-score) for UMAP analysis to generate a reduced-dimensionality matrix. Gene set enrichment analysis for both KEGG pathways and GO annotations was performed using the latest annotations from the KEGG REST API and org.Hs.eg.db R package (version 3.15.0), respectively. The clusterProfiler R package (version 4.4.4) mapped genes to these annotations, identifying statistically significant pathways and GO terms. Enrichment analyses considered a minimum gene set of 5, a maximum of 5000, and an FDR threshold of <0.05 as significant.

### 2.12 Statistical analysis

This study’s data normality was assessed using the Shapiro-Wilk test, and findings were presented as mean ± SEM, analyzed via GraphPad Prism 10. Appropriate statistical tests, including one-way or two-way ANOVA with Bonferroni corrections for normally distributed data adhering to homogeneity of variance, Welch’s ANOVA, or Dunnett’s T3 for unequal variances, and Kruskal-Wallis for non-normal distributions, were applied. Two-group differences were evaluated using two-tailed t-tests. Significance was established at p< 0.05.

## Results

### 3.1 Human and mouse brain transcriptomic atlases reveal developmental trajectory and cell subtype specific expression patterns of *PATJ*

To extend prior human genetic work implicating *PATJ* in metabolic disease and stroke and provide further insights into PATJ function, we examined existing large mouse and human brain single cell RNA sequencing atlases profiling *PATJ* expression across neural cell types and the lifespan [23–25]. Interestingly, during fetal development and infancy *PATJ* is particularly enriched and levels decrease into adulthood. This pattern of expression is seen in many other key genes involved in neurogenesis and vasculogenesis such as ephrin B1, paired box protein (*PAX*)*-6*, neuronal differentiation (*NeuroD*)-*1*, and beta-catenin (*CTNNB1*), among others (Figure 1A). Although *PATJ* is enriched in early life, its expression in a broad array of cells is still present into adulthood in human and mouse single cell RNA atlas data (Figure 1B-C). In particular, single cell RNA sequencing atlas of whole human or mouse brain demonstrates that *PATJ* expression is most prominent in cerebellar glutamatergic cells, but also highly expressed in endothelial and other brain vascular cells (particularly those of the choroid plexus; Figure 1B-C). Of note, the cell-specific distribution of other key genes previously suggested to interact with *PATJ* including *AMOT*, *PLEKHG5*, and *YAP1* demonstrate similar enrichment in cerebral vascular cell subtypes (see Supplementary Figure 1). Taken together, existing human and mouse cell type-specific gene expression data suggest functional roles for PATJ in both neuronal and vascular biology.

**Figure 1.**
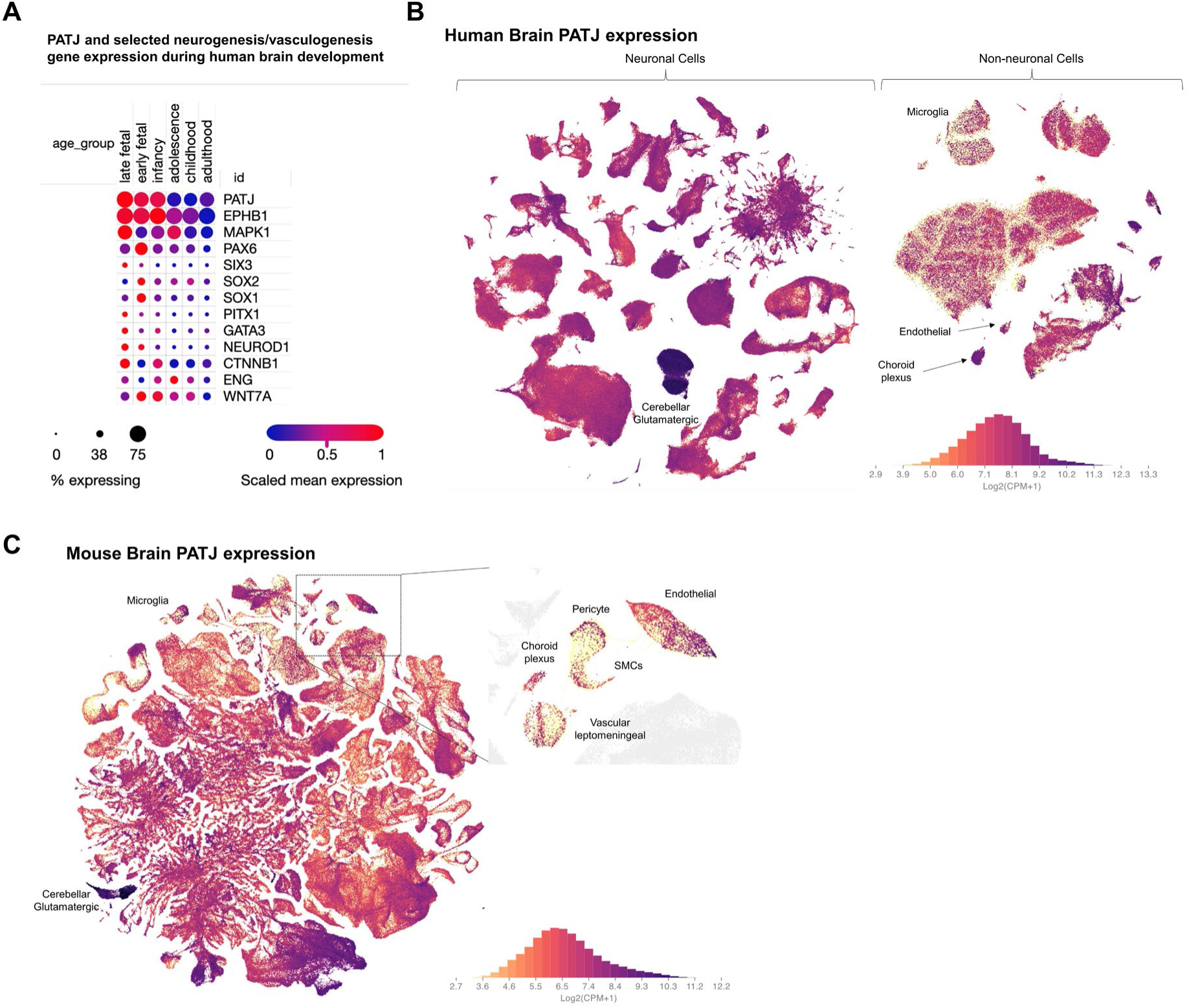
*PATJ* expression in human and mouse brain single cell RNA sequencing databases. (A) Dotplot demonstrating expression of *PATJ* and other key regulators of neurogenesis and vasculogenesis in the cortex. *PATJ* expression is highest during fetal development and infancy and decreases in adolescence and adulthood. Data from Zhu et al., 2023 (GSE204684) and accessed at singlecell.broadinstitute.org. (B) Human and (C) Mouse single cell RNA sequencing brain atlas UMAP data demonstrating ubiquitous expression of *PATJ* in adult brain cell subtypes, with inset highlighting vascular cell subtypes. Data from Siletti et al., 2023 and Yao et al., 2023, respectively and accessed at knowledge.brain-map.org.

### 3.2 Establishment of PATJ KO HEK Cell Line via CRISPR-Cas9

We designed two single-guide RNAs (sgRNAs) targeting exon 3 of the *PATJ* gene in Human Embryonic Kidney 293 (HEK 293) cells by CRISPR-Cas9. Using this strategy, we screened clonally edited populations by sequencing and identified two clonal lines with a frameshift mutation that effectively eliminated expression of the PATJ protein as well as one clonal line without editing to serve as an unedited control (Figure 2).

**Figure 2.**
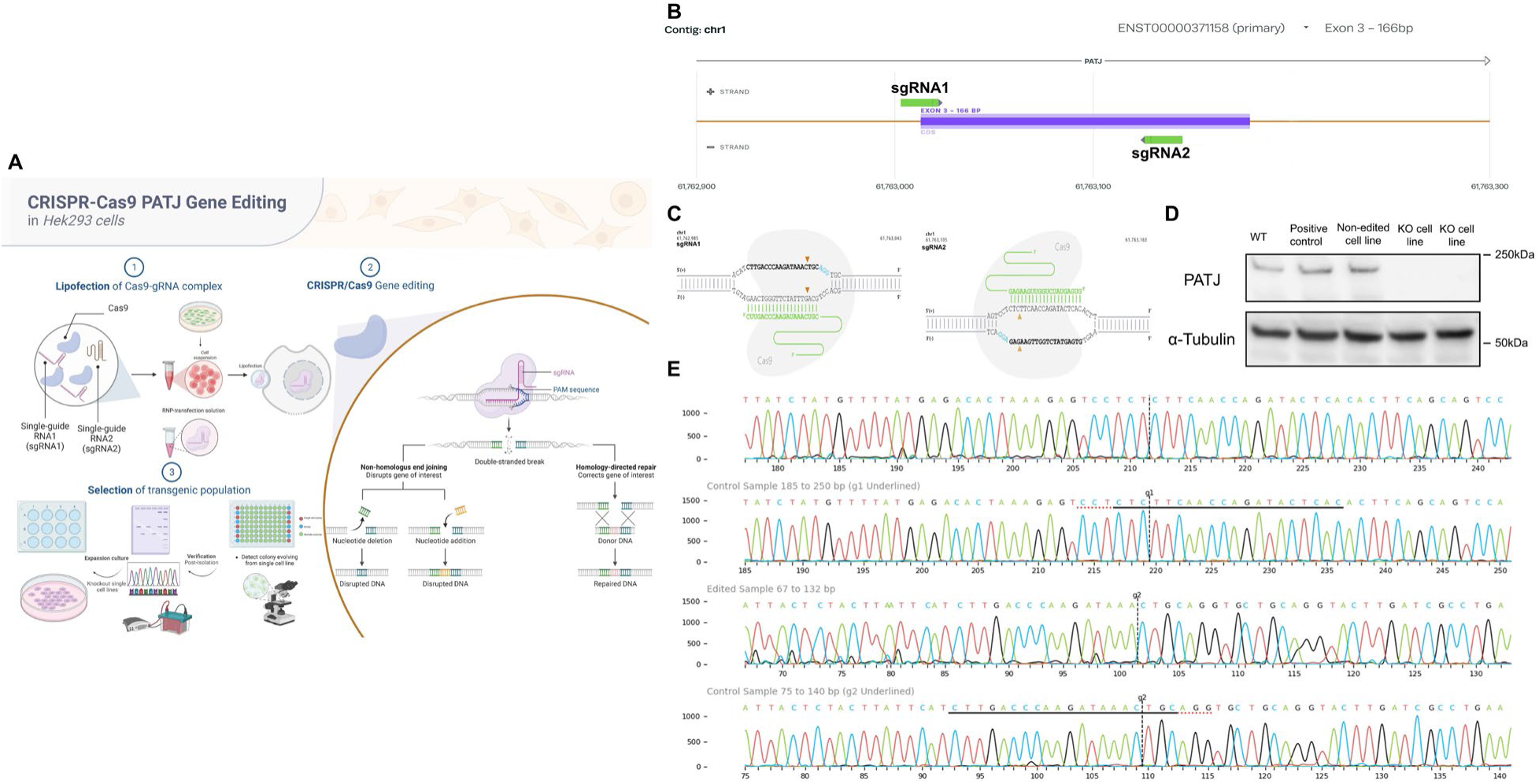
CRISPR-Cas9 mediated knock-out of *PATJ* in HEK293 cell lines and validation. (A) Schematic overview of the CRISPR-Cas9 gene editing workflow for *PATJ* in HEK293 cells. (B) Schematic representation of CRISPR-Cas9 strategy targeting exon 3 of the *PATJ* gene. Two sgRNAs were designed for precise gene editing. (C) Sequence alignment for sgRNA1 and sgRNA2 targeting specific sites within exon 3 of the *PATJ* gene. (D)Western blot analysis verifying the absence of PATJ protein expression in knockout cell lines, establishing the efficacy of gene disruption as compared to the WT control. (E) Sanger sequencing showing the insertion, leading to a frameshift mutation in the *PATJ* gene of KO cells.

### 3.3 PATJ modulates stress responses in mammalian cells and *C. elegans*

In mammalian cells, we found that PATJ deficiency enhanced vulnerability to mitochondrial stress. A statistically significant discrepancy in growth was noted between PATJ KO and WT cells (Fig. 3A). This variation suggests that the absence of PATJ may inherently limit the proliferation of cells. Rotenone, a mitochondrial complex I inhibitor, was used to model energetic stress in PATJ-deficient contexts. Rotenone treatment markedly impaired PATJ KO cell survival after 72 hours compared to wildtype HEK293 cells (Fig. 3B). Interestingly, differences in oxygen consumption were also observed in vitro following administration of oligomycin and rotenone/antimycin (Fig. 3C) while preserving spare respiratory capacity. The sensitivity to oligomycin, an inhibitor of ATP synthesis, and rotenone/antimycin, inhibitors of complex I and complex III, suggests a greater dependence on oxidative phosphorylation in PATJ KO cells. This data collectively indicates that PATJ may contribute to cell survival and adaptability in conditions of metabolic stress.

**Figure 3.**
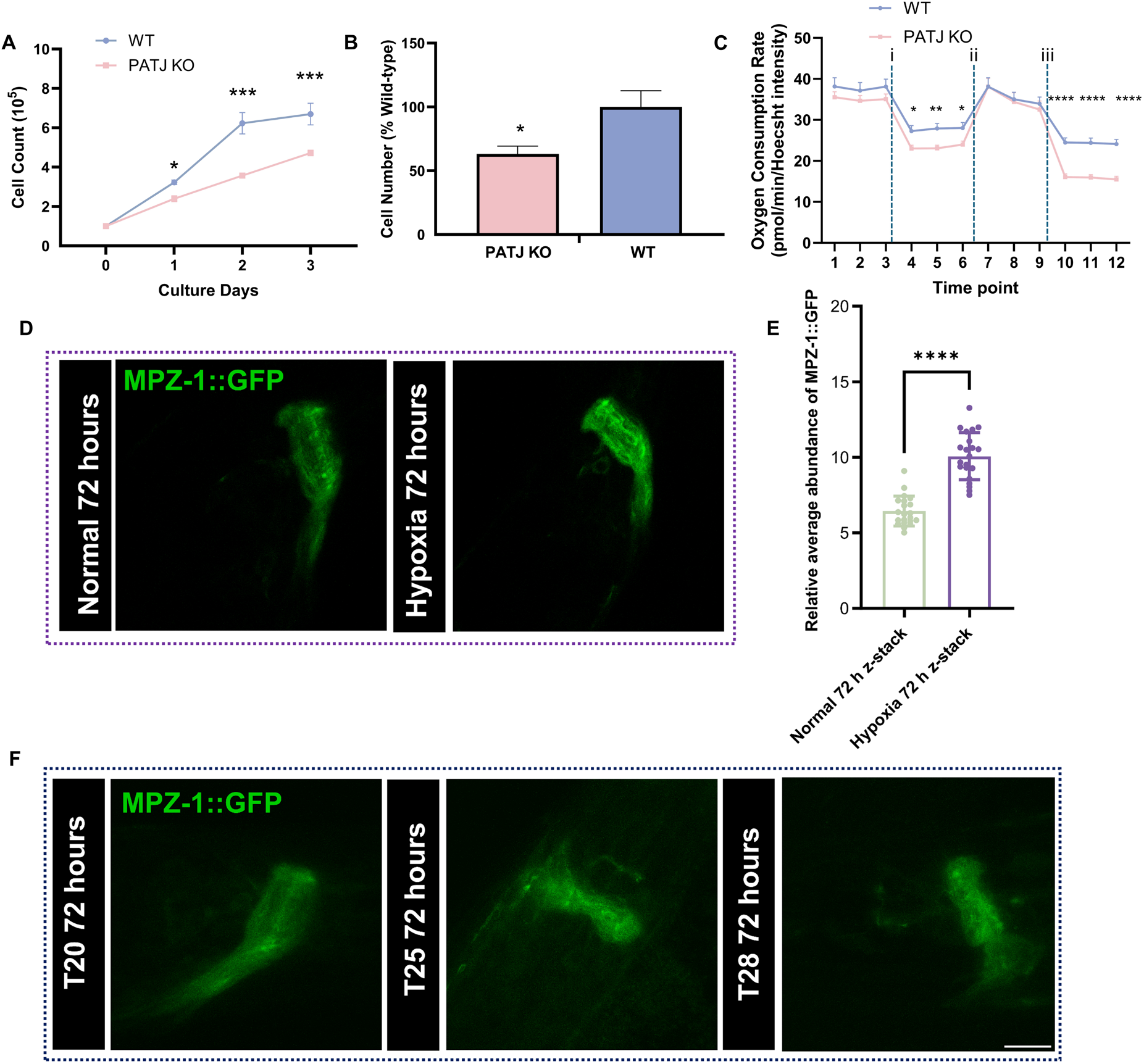
PATJ modulates stress responses in mammalian cells and *C. elegans*. (A) Growth comparison of PATJ knockout (KO) and wild-type (WT) HEK293 cells. (B) Survival rates of PATJ KO and WT cells after 72 hours of treatment with rotenone. (C) Oxygen consumption rates (OCR) of PATJ KO and WT cells following treatment with oligomycin (i), FCCP (ii), and antimycin A/rotenone (iii). n=4-6 for each group. (D) Representative confocal images of endogenous MPZ-1::GFP expression in *C. elegans* under hypoxia (0.1% O2) or normoxia for 72 hours. Scale bars: 10 µm. (E) Quantification of MPZ-1::GFP fluorescence intensity under hypoxic and normoxic conditions. (F) Representative confocal images of MPZ-1::GFP expression in *C. elegans* under heat stress at 20°C, 25°C, or 28°C for 72 hours. Scale bars: 10 µm. n>20 animals per condition for *C. elegans* experiments. * p<0.05, ** p<0.01, *** p<0.001, **** p<0.0001.

To investigate the evolutionary conservation of PATJ’s role in stress responses, we examined the expression of MPZ-1, a *C. elegans* ortholog of PATJ, under hypoxic and heat stress conditions using a CRISPR-generated MPZ-1::GFP reporter strain. Remarkably, exposure to hypoxia (0.1% O2) for 72 hours resulted in a significant upregulation of MPZ-1::GFP expression compared to normoxic controls (Fig. 3D,E). In contrast, heat stress at 25°C or 28°C for 72 hours did not significantly alter MPZ-1::GFP levels relative to the 20°C control (Fig. 3F). These findings suggest that the induction of PATJ expression in response to hypoxia is specific and conserved in C. elegans.

### 3.4 Phenotypic Characterization Reveals Reduced Migration in PATJ KO Cells Ameliorated by Overexpression

Prior work has implicated PATJ in developmental processes and suggested interaction with cytoskeletal proteins[26]. As such we hypothesized that cell migration may be affected in *PATJ* KO cells as assessed by an *in vitro* scratch assay. In this assay a mechanical scratch is introduced into cells grown at high confluence and the resulting ‘scar’ monitored for cell migration and closure. Cell proliferation is inhibited with the addition of mitomycin C. We found a notable reduction in cellular migration in *PATJ* KO cells compared to both WT HEK293, while accelerated migration was seen in cells overexpressing PATJ (Figure 4). Quantitative analysis demonstrated a significant increase in the gap width of *PATJ* KO cells at 24 hours, which persisted to 48 hours post-scratch. The migration impairment observed in *PATJ* KO cells was effectively rescued by overexpression of PATJ to levels comparable to WT cells. These results indicate a critical role for PATJ in cell migration following an injury.

**Figure 4.**
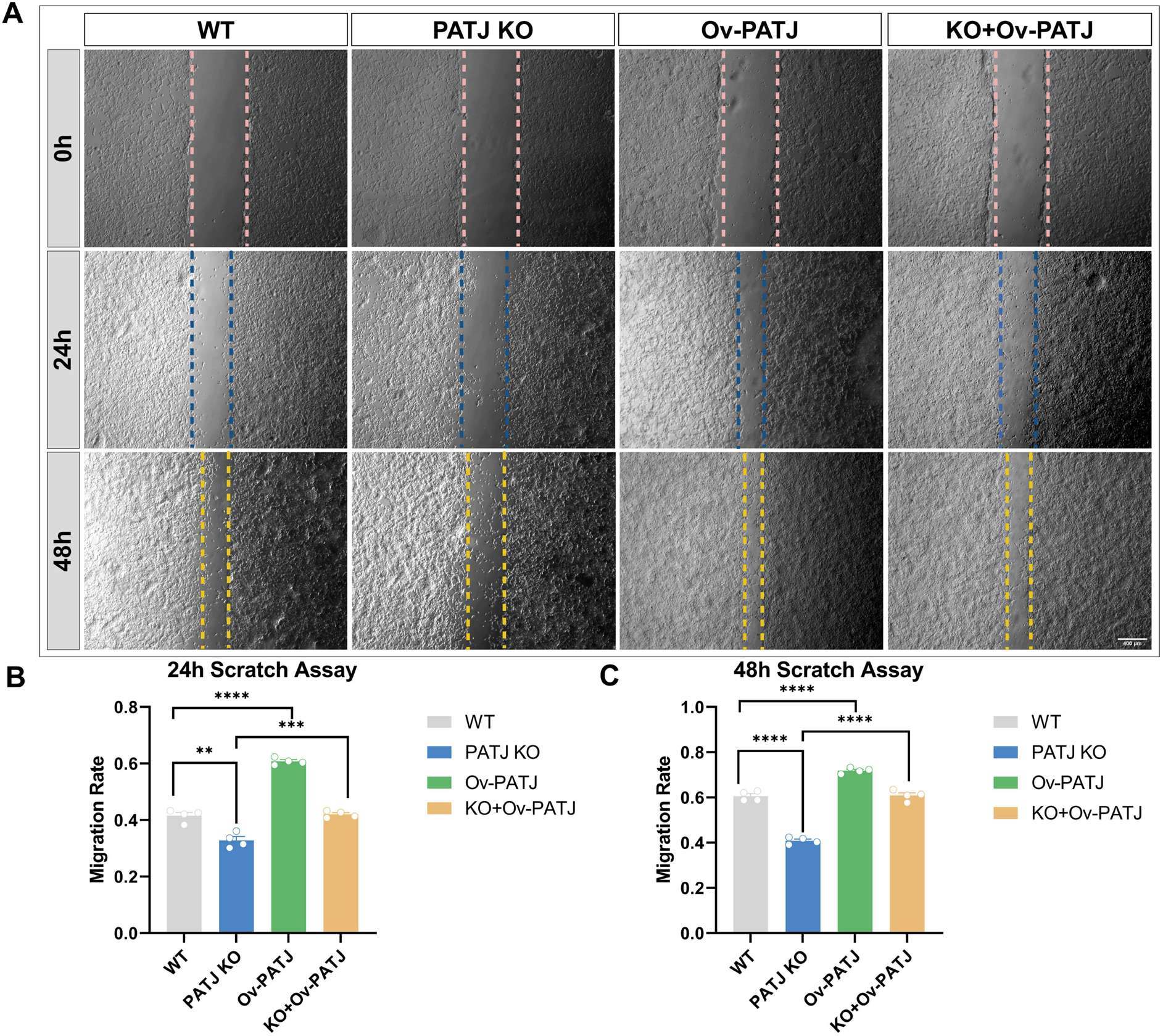
Comparative analysis of cell migration in *PATJ*-modified HEK293 Cells. (A) Scratch assay time-lapse images at 0, 24, and 48 hours post-wounding display cell migration for various HEK293 cell lines: wild-type (WT), *PATJ* knock-out (KO), PATJ overexpression (Ov-PATJ), and KO with PATJ overexpression (KO+Ov-PATJ). (B) Quantitative analysis of cell migration at 24 hours post-scratch. (C) Quantitative analysis of cell migration at 48 hours post-scratch. n=4 for each group. * p<0.05, ** p<0.01, *** p<0.001, **** p<0.0001.

### 3.5 Regulatory interplay between PATJ deletion and YAP1 subcellular dynamics

To further investigate the underlying mechanisms of PATJ-mediated cellular regulation, we delineated the influence of *PATJ* deletion on the regulation of YAP1, a key effector within the Hippo signaling pathway. Western blot analysis demonstrated an upregulation in YAP1 protein levels in *PATJ* KO cells compared to WT (Figure 5A), suggesting a regulatory interplay between PATJ and YAP1 expression or stability.

**Figure 5.**
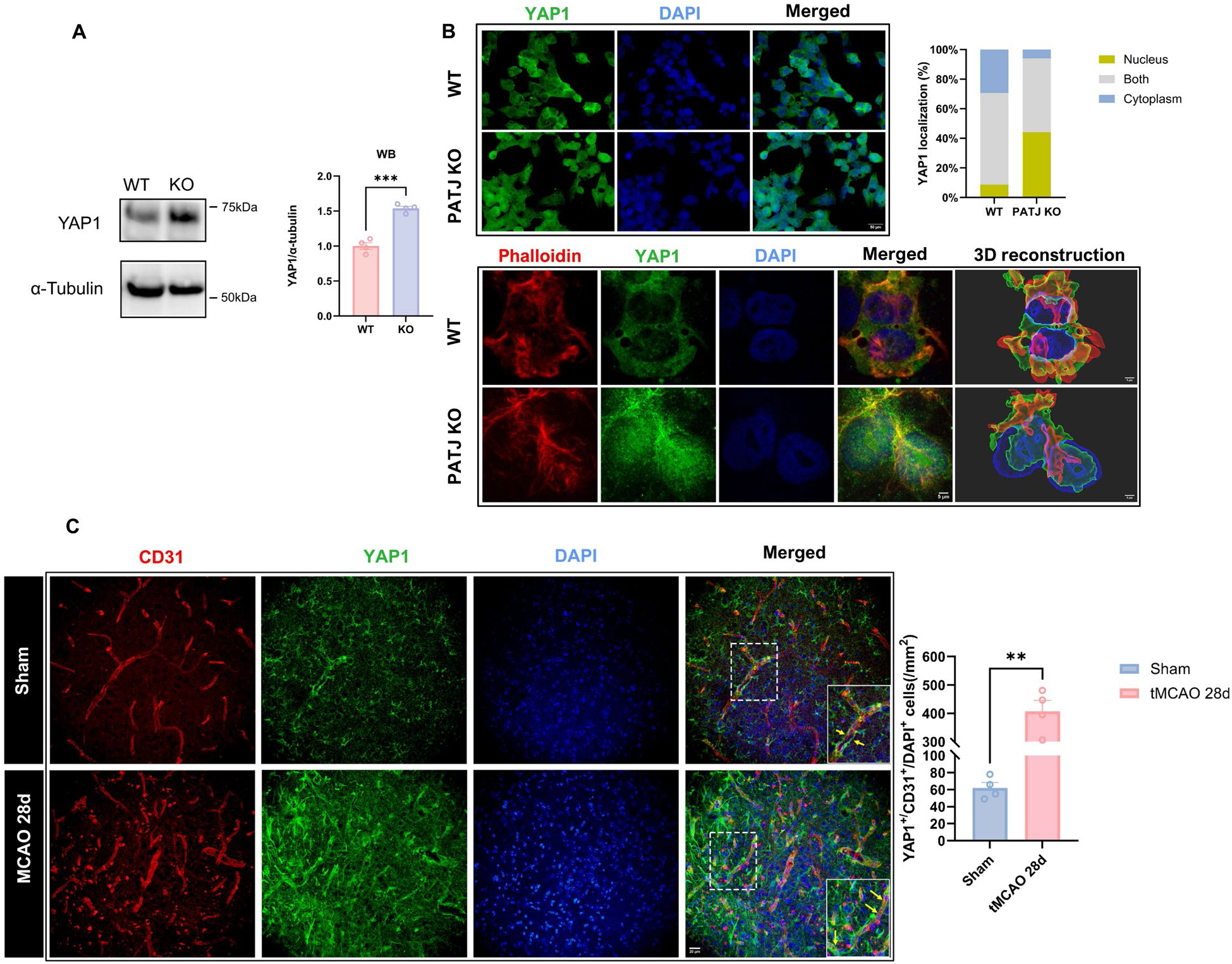
YAP1 expression and localization in PATJ-modified HEK293 cells and ischemic stroke. (A) Western blot analysis of YAP1 protein levels in WT and *PATJ* KO cells, with quantification showing a significant decrease in KO cells. (B) Immunofluorescence staining of YAP1 (green) and Phalloidin (red) in WT and *PATJ* KO cells, with DAPI staining of nuclei (blue). The right panel shows the percentage of YAP1 localization in the nucleus, cytoplasm, or both. 3D reconstruction of the staining pattern is also presented. (C) Representative images of endothelial cell marker CD31 (red) and YAP1 (green) in brain tissue from sham and 28 days post-middle cerebral artery occlusion (MCAO) mice, with DAPI staining of nuclei (blue) and merged images highlighting co-localization. n=4 for each group. *** p<0.001.

Interestingly, although YAP1 was upregulated in the *PATJ* KO cells, KO cells demonstrated immunofluorescence imaging revealed YAP1 accumulation in the nucleus, contrasting with the primarily cytoplasmic presence in WT cells (Figure 5B). Quantitative analysis confirmed a significant increase in the percentage of cells displaying predominantly nuclear YAP1 localization in the absence of PATJ. The nuclear enrichment of YAP1 in the absence of PATJ implies a potential reprogramming of YAP1-dependent transcriptional activity. The observed changes in YAP1 localization and expression in *PATJ* KO cells confirm the role of PATJ in influencing key cellular functions and suggest a possible interaction with YAP1’s activity within cells.

To extend these findings to an in vivo context, we examined YAP1 expression and localization in a mouse model of ischemic stroke. Immunohistochemical analysis of brain sections from mice subjected to tMCAO revealed a distinct alteration in YAP1 subcellular localization in endothelial cells 28 days post-stroke compared to sham-operated controls (Figure 5C). In sham group, endothelial YAP1 expression was primarily localized to the cell surface. In contrast, tMCAO mice exhibited a marked increase in nuclear YAP1 within endothelial cells in the peri-infarct region, as evidenced by the co-localization of YAP1, CD31, and the nuclear marker DAPI. Quantitative analysis of cells positive for all three markers confirmed a significantly higher number of endothelial cells with nuclear YAP1 in tMCAO mice compared to sham controls. These findings suggest that ischemic stroke promotes the nuclear translocation of YAP1 in endothelial cells, potentially as a vascular adaptive response to ischemic injury.

### 3.6 Temporal profiling of PATJ expression in endothelial cells following stroke

To further explore the changes in PATJ after ischemic stroke, we characterized the temporal patterns of PATJ expression following transient middle cerebral artery occlusion (tMCAO) in mice (Figure 6). We observed a reduction in PATJ expression within the endothelial cells in the penumbral regions. This decrease was statistically significant when compared to the sham-operated controls, suggesting that PATJ may be acutely sensitive to ischemic conditions. As the post-stroke timeline progressed, a gradual yet significant increase in PATJ expression became evident. As the recovery phase progressed, the colocalization of PATJ within the peri-ischemic endothelium gradually increased. By day 28 post-tMCAO, not only did PATJ expression surpass its original levels, but its colocalization with endothelial markers also significantly increased beyond the baseline observed in sham-operated controls. This enhanced colocalization in the later stages post-stroke signifies a possible adaptive augmentation of PATJ in post-stroke endothelial response to drive vascular remodeling and repair.

**Figure 6.**
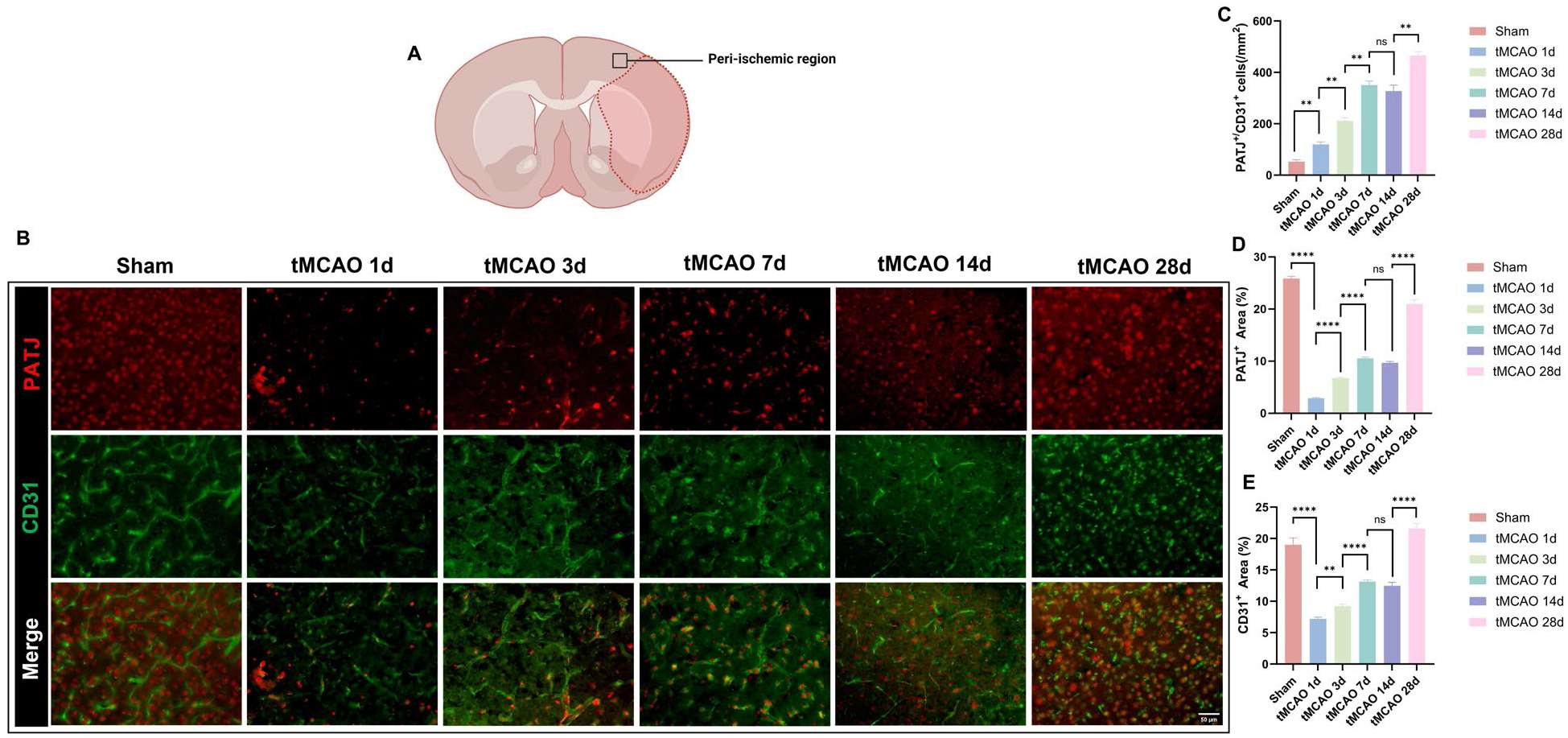
Temporal expression of Patj in endothelial cells post-tMCAO. (A)Schematic representation of the peri-ischemic region observed in the tMCAO mouse model. (B) Representative immunofluorescence images showing double staining of Patj (red) and CD31 (green) in brain sections from sham and tMCAO-treated mice at days 1, 3, 7, 14, and 28 post-stroke. (C) Quantitative analysis of Patj/CD31 double positive cells per mm² in the peri-ischemic region over time (D) Quantitative analysis of Patj positive cells in the peri-ischemic area over time. (E) Quantitative analysis of CD31 positive signal dynamics in the peri-ischemic zone. n=4 for each group. ** p<0.01, **** p<0.0001.

### 3.7 Temporal shifts in PATJ localization within neurons following stroke

Building on our observations of PATJ expression in endothelial cells following stroke, we extended our analysis to examine its neuronal distribution under physiological conditions and after ischemic injury. In the neuronal domain, PATJ exhibited a distinct expression profile in response to ischemic conditions (Figure 7). Contrary to the upregulation observed in endothelial cells, neuronal PATJ demonstrated a marked decrease following ischemic stroke. This decrement persisted even 28 days post-stroke, suggesting a sustained impact of the ischemic event on neuronal PATJ presence. Neuronal expression of PATJ in hemisphere contralateral to stroke was unaffected. This persistent decrease in neuronal PATJ expression contrasts sharply with the temporal enhancement seen in endothelial cells, underscoring a possible divergence in PATJ function between these cell types after stroke.

**Figure 7.**
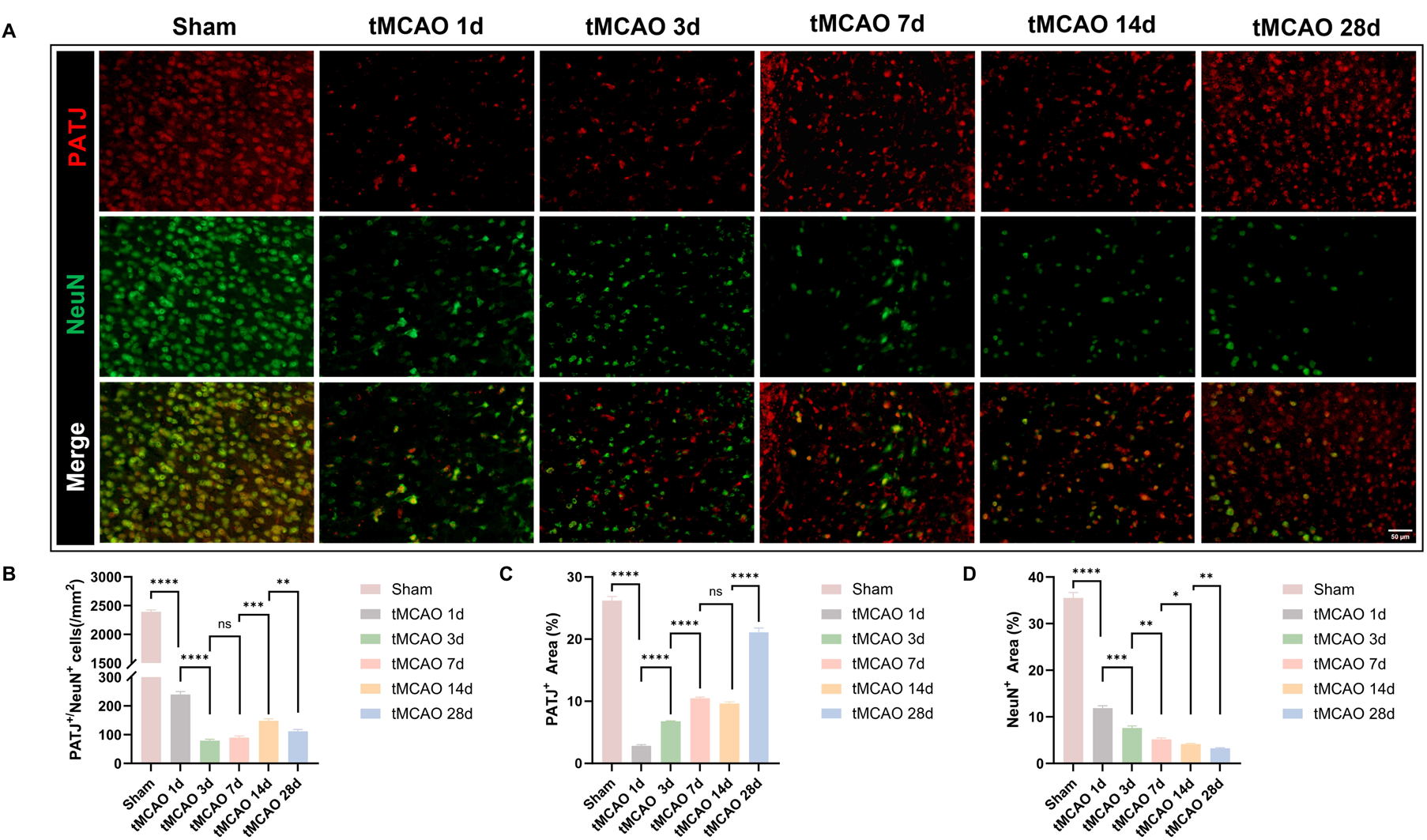
Temporal shifts in *Patj* expression in neurons after tMCAO. (A) Representative immunofluorescence images showing double staining of Patj (red) and the neuronal marker NeuN (green) in brain sections from sham and tMCAO mice at 1, 3, 7, 14, and 28 days post-stroke. (B) Quantitative analysis of Patj/NeuN double-positive cells per mm² in the peri-ischemic region across the specified time points. (C) Quantitative analysis of Patj positive signal in the peri-ischemic region over time (D) Quantitative analysis of NeuN positive cells in the peri-ischemic zone at each time point post-tMCAO. n=4 for each group. * p<0.05, ** p<0.01, or *** p<0.001, **** p<0.0001.

### 3.8 Transcriptomic profiling reveals PATJ’s influence on vascular and cellular processes

We conducted RNA-sequencing on *PATJ* KO, *PATJ* KO with PATJ overexpression (KO+Ov), and WT cell lines, revealing distinct transcriptional landscapes. Principal component analysis (PCA) validated our approach, showing clear group segregation (Figure 8A). A core set of 112 genes were differentially regulated across the three cell types, suggesting essential elements of the PATJ-dependent transcriptional program (Figure 8B). Runt-related transcription factor 1 (RUNX1) is critically involved in hematopoiesis and has been implicated in endothelial functions[27], suggesting a potential link between PATJ modulation and vascular remodeling, a process vital in post-stroke recovery. Hairy/enhancer-of-split related with YRPW motif 1 (*HEY1*), a transcriptional repressor regulated by the Notch signaling pathway, is known to influence vascular development and integrity[28], aligning with the observed changes in PATJ-mediated gene expression and pointing toward a role in angiogenesis and possibly neurogenesis. Nuclear protein 1 (*NUPR1*), involved in stress responses and cellular adaptation[29], may offer insights into the cellular mechanisms by which PATJ contributes to the resilience and migration of cells in response to ischemic conditions induced by stroke. Furthermore, Hexokinase 2 (*HK2*), central to the glycolytic pathway, demonstrated markedly reduced expression levels in PATJ KO cells suggesting a greater dependence on oxidative phosphorylation and supporting the Seahorse assay results also showing a shift in metabolic requirements during cellular stress[30]. Heatmap and volcano plot analyses further highlighted significant upregulation and downregulation of genes associated with immune response, growth regulation, and signal transduction, indicating PATJ’s diverse regulatory impact (Figures 8C, E, G, and 8D, F, H). Gene Ontology (GO) and KEGG pathway analyses connected the DEGs to crucial stroke-related processes in the KO+Ov group, including vascular development and response to hypoxia, with pathways like ‘p53 signaling’ and ‘Notch signaling’ also being enriched (Figure 8I,J).

**Figure 8.**
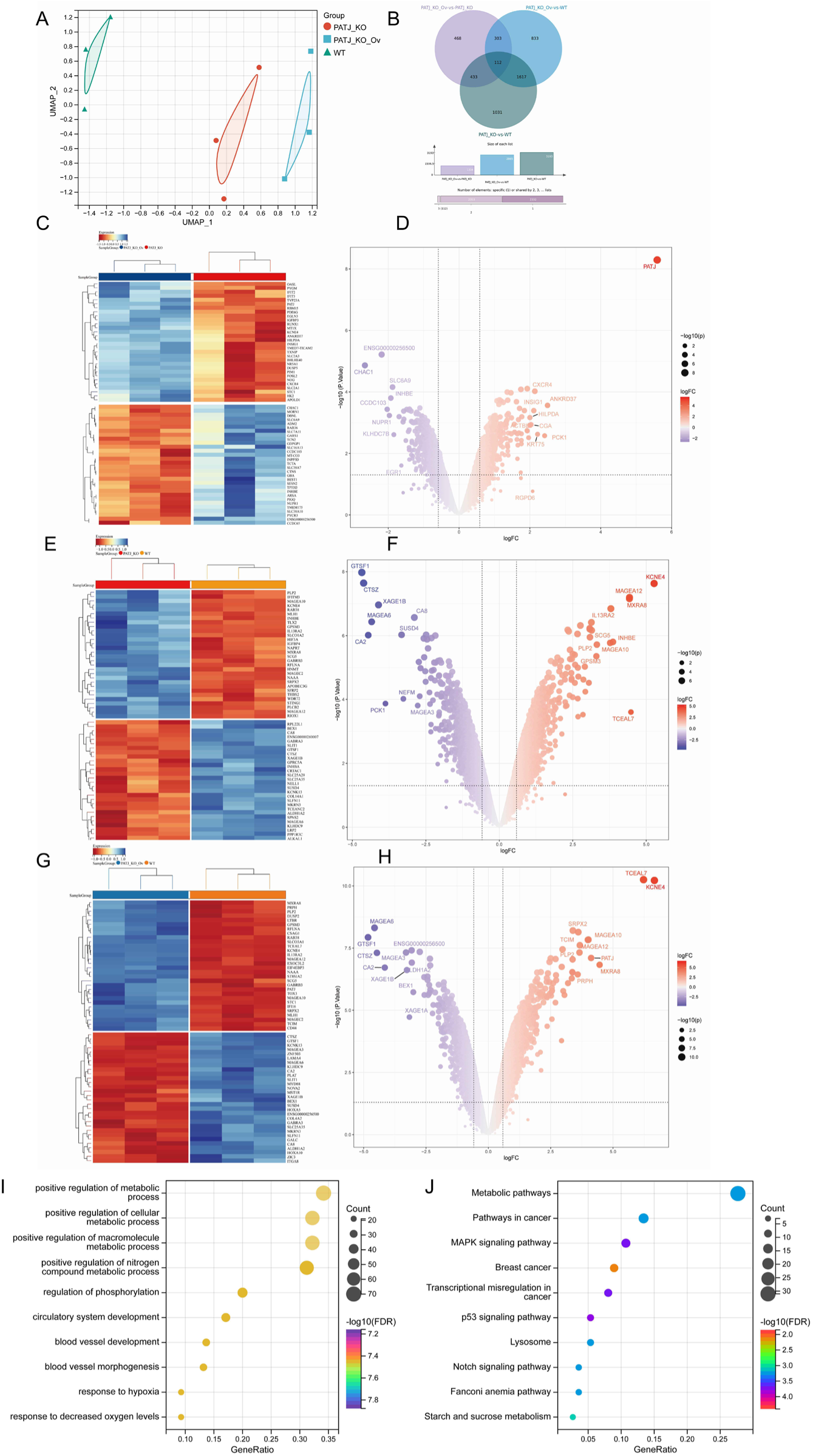
Transcriptomic landscape and pathway analysis in *PATJ*-modulated HEK293 cells. (**A)** UMAP visualization segregating gene expression profiles of wild-type (WT), *PATJ* knock-out (KO), and *PATJ* knock-out with overexpression (KO+Ov) HEK293 cells. (**B)** Venn diagram highlighting the shared and unique differentially expressed genes among WT, KO, and KO+Ov cell groups. (**C, E, G)** Heatmaps of gene expression contrasts for KO+Ov vs. KO, KO vs. WT, and KO+Ov vs. WT, respectively, with color intensity denoting gene expression levels. (**D, F, H)** Volcano plots detailing significant gene expression changes for KO+Ov vs. KO, KO vs. WT, and KO+Ov vs. WT, with upregulated genes in red and downregulated genes in blue. (**I)** Gene Ontology (GO) enrichment analysis bubble chart for the KO+Ov vs. KO comparison. (**J)** Kyoto Encyclopedia of Genes and Genomes (KEGG) pathway enrichment analysis bubble chart for the KO+Ov vs. KO comparison.

## Discussion

Ischemic stroke results in significant vascular and neuronal damage. Cerebral endothelial cells (ECs) ischemic damage increases vascular permeability and disrupts the blood-brain barrier[31–33]. Thus, optimizing the beneficial effects of ECs during stroke recovery is an attractive target to improve stroke outcome[34, 35]. PATJ is known for its involvement in tight junction assembly and maintenance of epithelial cell polarity[8–11, 36–40]. Although GWAS study has linked it as a significant genetic locus mediating stroke functional outcome[13], and only recently has preclinical evidence suggested its role in ECs function following stroke and other biological processes [16].

In our study, we investigated PATJ’s multifaceted role in cell physiological conditions and extending into pathological scenarios via an animal stroke model. We successfully employed CRISPR-Cas9 technology for the establishment of a *PATJ* KO in HEK293 cell lines, which revealed roles for in cell migration, response to mitochondrial stress, and the regulatory dynamics with YAP1. These insights underscore PATJ’s critical role in cellular adaptability and survival mechanisms and provide a starting point for further mechanistic study.

Our findings reveal a significant reduction in cellular migration in *PATJ* KO HEK293 cells, which can be rescued by inducing the overexpression of PATJ. This aligns with previous studies highlighting PATJ’s involvement in regulating directional migration of mammalian epithelial cells and interaction with cytoskeletal proteins [9, 26]. Our data also reveal an enhanced vulnerability to mitochondrial stress in PATJ-deficient cells with reduced proliferation in rotenone and altered oxygen consumption following oligomycin, highlighting the gene’s contribution to cellular resilience against metabolic challenges. This aspect is particularly relevant in the context of stroke, where cellular energy demands are profoundly altered. The ability of cells to withstand and adapt to such conditions is critical for survival and recovery, suggesting that interventions aimed at modulating PATJ expression or its targets could improve cellular resilience in ischemic injury.

The regulatory interplay between PATJ and YAP1, especially under conditions of *PATJ* depletion, introduces a new angle for further study in understanding EC response mechanisms to ischemic stroke, especially considering YAP1 is highly enriched in brain vascular cells (see Supplementary Figure 1;[24]). The observed changes suggest that *PATJ* deletion may trigger YAP1 signaling compensatory mechanisms. Emerging evidence suggests that nuclear translocation of the transcription factor, YAP1 modulates the expression of genes involved in angiogenesis[41–53]. Intriguingly, our in vivo studies corroborate and extend these findings. We observed a marked increase in nuclear YAP1 within endothelial cells in the peri-infarct region, suggesting a potential vascular response to ischemic injury. While the precise role of YAP1 in post-stroke angiogenesis remains to be fully elucidated, this nuclear translocation may indicate its involvement in vascular adaptation processes. Angiogenesis, which ECs are principally involved, begins in the zones surrounding the ischemic event within the first 12 hours and can persist for more than three weeks[54, 55]. Angiogenesis is necessary for the repair and remodeling of vascular structures and also promotes neural plasticity, as evidenced in both clinical and preclinical stroke studies[56]. The increased nuclear localization of YAP1 in endothelial cells of the peri-infarct region could represent part of a complex adaptive response, potentially influencing vascular remodeling in the recovering brain tissue. Thus further understanding the mechanistic connections between PATJ and YAP1 may aid in uncovering new therapeutics to enhance post-stroke angiogenesis and tissue repair.

The dynamic regulation of PATJ expression in ECs following ischemic stroke, characterized by an initial decrease and subsequent increase, suggests a tightly controlled response to ischemic injury. This biphasic pattern might reflect the role of PATJ in interacting with EC repair processes post-stroke. Interestingly, our findings in *C. elegans* demonstrate that the specific upregulation of MPZ-1, the *C. elegans* ortholog of PATJ, under hypoxic conditions is particularly relevant to understanding PATJ’s potential role in ischemic stroke adaptation. As ischemic stroke is characterized by reduced blood flow leading to hypoxia and metabolic stress in the affected brain regions, the observation that PATJ expression is induced by hypoxia in *C. elegans* suggests that this response may be an evolutionarily conserved mechanism to cope with oxygen deprivation.

The increased sensitivity of PATJ-deficient mammalian cells to mitochondrial insults further supports the notion that PATJ plays a crucial role in cellular adaptation to metabolic stress. Given that ischemic stroke triggers hypoxia and metabolic dysfunction, the evolutionarily conserved ability of PATJ to orchestrate adaptive responses may have important implications for understanding its influence on stroke outcomes.Future work will be necessary to understand whether modulating PATJ expression also modulates vascular remodeling to enhance stroke recovery. Conversely, the sustained decrease in neuronal PATJ expression following stroke further emphasizes the need for additional research in understanding the cell-type-specific regulation of PATJ to enhance stroke recovery.

Our transcriptomic analysis, centered on PATJ’s modulation, illuminates its pivotal role in regulating genes and pathways critical for vascular remodeling and recovery post-stroke. Notably, the downregulation of *RUNX1* and *HEY1* in *PATJ* KO cells, implicated in endothelial functions and angiogenesis[27, 28, 57], underscores PATJ’s influence on vascular development. This, coupled with NUPR1’s involvement in cellular stress responses and HK2’s role in metabolic adaptation[29, 30], highlights metabolic influences regulated by PATJ. Notably, the upregulation of noggin (*NOG*) with PATJ overexpression and its down-regulation in the PATJ KO, suggests a previously unexplored connection between PATJ and this key regulator of neural development and neurogenesis [58]. Similarly, *RBM15* stands out with its substantial upregulation in the KO+Ov cell line and downregulation in the PATJ KO. RBM15 is integral to RNA metabolic processes, influencing not only mRNA splicing but also chromatin organization and gene expression of pathways critical for cellular identity and function. RBM15 dysregulation along with enrichment of the Notch signaling pathway in KEGG analysis aligns with our findings implicating PATJ in the modulation of vascular-related functions [59]. Interestingly, the chemokine C-X-C motif receptor (*CXCR*)-*4-* was also downregulated in the PATJ KO cell line, and is essential for chemotaxis and hematopoietic cell lineages [60] as well as cooperates with Notch signaling to facilitate arterial differentiation [57]. Taken together, these findings support a role for PATJ in influencing cell migration and differentiation. In contrast, genes such as *CHAC1* (involved in glutathione metabolism and apoptosis), *SLC6A9* (a neurotransmitter transporter), and *INHBE* (a member of the TGF-beta superfamily) exhibited considerable downregulation in KO+Ov cells and upregulation in *PATJ* KO cells. These changes may reflect PATJ’s involvement in the regulation of redox homeostasis, neurotransmission, and cell growth signaling pathways. The deregulation of these genes with genetic manipulation of PATJ expression coupled with the metabolic and cell biological phenotypes demonstrated in this study could suggest possible mechanistic targets for PATJ to exert effects on cellular stress responses, synaptic signaling, and tissue homeostasis. These findings, aligned with the validated GO and KEGG pathway functions, suggest PATJ’s integral role in orchestrating cellular resilience, angiogenesis, and neurogenesis through pathways like Notch signaling. Future research should focus on dissecting these pathways to unlock new therapeutic avenues for stroke recovery, emphasizing the need for targeted interventions that leverage PATJ’s regulatory capacity.

Extending the foundational work by Medina-Dols et al. on PATJ’s role in modulating endothelial to mesenchymal transition (EndMT) through key pathways [16], our investigation provides additional insights and corroborates key findings with complementary techniques. Employing CRISPR-Cas9 technology, we illuminate PATJ’s broader influence on cellular dynamics and vascular remodeling. Our findings reveal significant PATJ-driven alterations in cellular migration and mitochondrial stress response, highlighting a regulatory interplay with the YAP1 signaling pathway, particularly highlighting YAP1’s nuclear translocation—a phenomenon also partially supported by Medina-Dols and colleagues findings. Our RNA-seq analysis reveals extensive transcriptional reprogramming under PATJ modulation, identifying dysregulation in genes critical to vascular development, stress response, and metabolism. Building upon Medina-Dols et al.’s observations of sustained PATJ downregulation[16], our in vivo analysis showcases a dynamic, time-dependent expression pattern in endothelial cells, suggesting PATJ’s integral role in both the immediate response and subsequent recovery processes. This biphasic expression underscores the complexity of PATJ’s involvement beyond EndMT, implicating it in essential pathways for vascular development and recovery. Thus, this work significantly extends our understanding of PATJ’s role in stroke pathophysiology, and supports further work in this pathway to pave new avenues for targeted therapeutic interventions in stroke rehabilitation and vascular repair.

In conclusion, our research advances the understanding of PATJ’s multifaceted roles in cellular and vascular physiology, particularly its evolutionarily conserved function in modulating cellular responses to hypoxic stress, and also highlights the potential of targeting PATJ-mediated molecular network for therapeutic intervention in stroke recovery. Future studies will explore the detailed mechanisms by which PATJ influences EC function and vascular remodeling in stroke recovery other pathologic states, further illuminating its role in disease modulation and therapeutic potential.

---

**Supplementary Figure 1.**
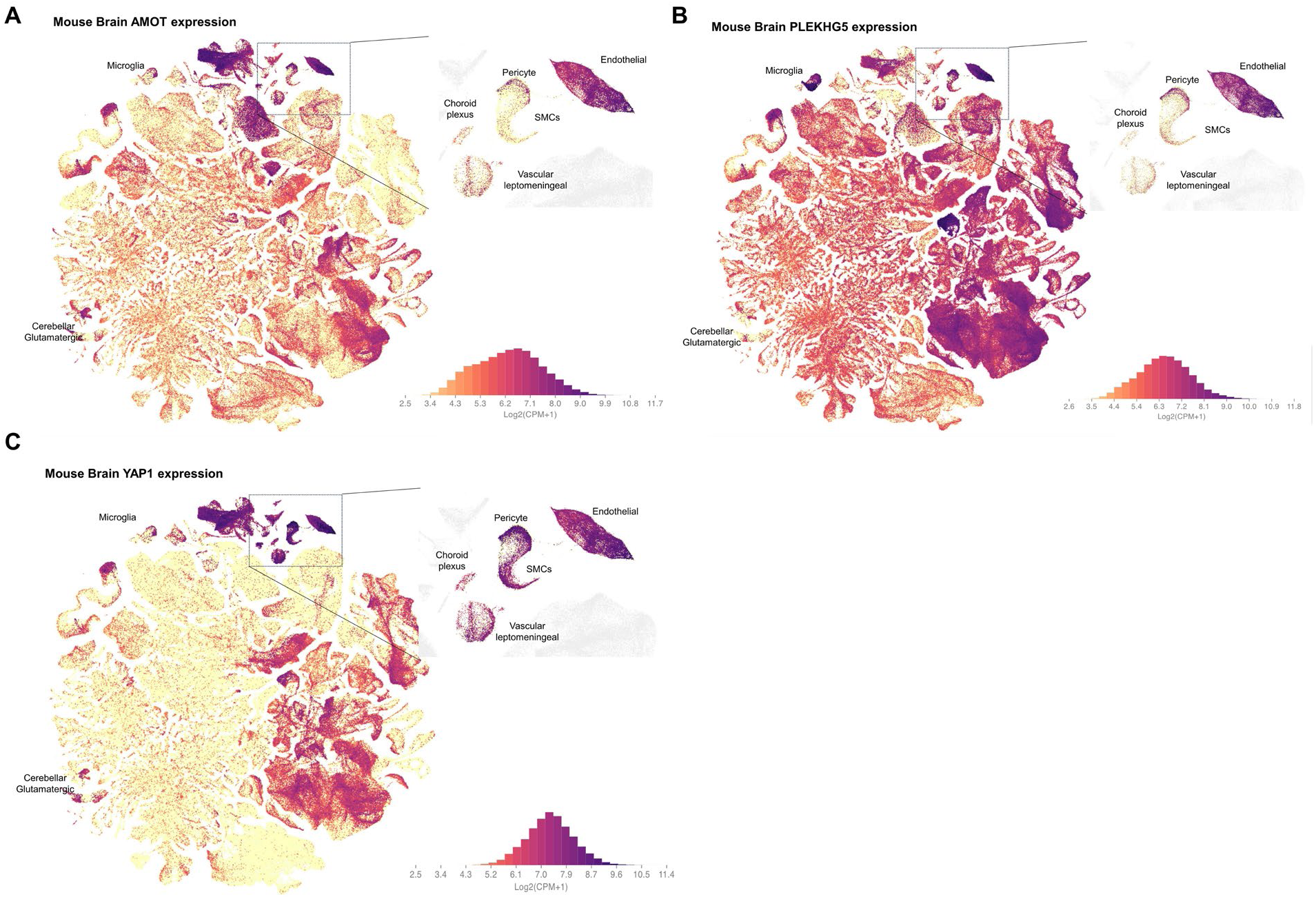
Mouse single cell RNA sequencing brain atlas UMAP data demonstrating cell subtype expression of (A) *Amot* (B) *Plekhg5*, and (C) *Yap1* in adult whole mouse brain, with inset highlighting vascular cell subtypes. Data from Yao et al., 2023 and accessed at knowledge.brain-map.org.

## References

1. Nogueira RG, Jadhav AP, Haussen DC, Bonafe A, Budzik RF, Bhuva P, Yavagal DR, Ribo M, Cognard C, Hanel RA et al: Thrombectomy 6 to 24 Hours after Stroke with a Mismatch between Deficit and Infarct. N Engl J Med 2018, 378(1):11–21.

2. George PM, Steinberg GK: Novel Stroke Therapeutics: Unraveling Stroke Pathophysiology and Its Impact on Clinical Treatments. Neuron 2015, 87(2):297–309.

3. Chung AS, Ferrara N: Developmental and pathological angiogenesis. Annu Rev Cell Dev Biol 2011, 27:563–584.

4. Marti HJ, Bernaudin M, Bellail A, Schoch H, Euler M, Petit E, Risau W: Hypoxia-induced vascular endothelial growth factor expression precedes neovascularization after cerebral ischemia. Am J Pathol 2000, 156(3):965–976.

5. Andjelkovic AV, Xiang J, Stamatovic SM, Hua Y, Xi G, Wang MM, Keep RF: Endothelial Targets in Stroke: Translating Animal Models to Human. Arterioscler Thromb Vasc Biol 2019, 39(11):2240–2247.

6. Liu J, Wang Y, Akamatsu Y, Lee CC, Stetler RA, Lawton MT, Yang GY: Vascular remodeling after ischemic stroke: mechanisms and therapeutic potentials. Prog Neurobiol 2014, 115:138–156.

7. Krupinski J, Kaluza J, Kumar P, Kumar S, Wang JM: Role of angiogenesis in patients with cerebral ischemic stroke. Stroke 1994, 25(9):1794–1798.

8. Tan B, Yatim S, Peng S, Gunaratne J, Hunziker W, Ludwig A: The Mammalian Crumbs Complex Defines a Distinct Polarity Domain Apical of Epithelial Tight Junctions. Curr Biol 2020, 30(14):2791–2804 e2796.

9. Shin K, Straight S, Margolis B: PATJ regulates tight junction formation and polarity in mammalian epithelial cells. J Cell Biol 2005, 168(5):705–711.

10. Penalva C, Mirouse V: Tissue-specific function of Patj in regulating the Crumbs complex and epithelial polarity. Development 2012, 139(24):4549–4554.

11. Michel D, Arsanto JP, Massey-Harroche D, Beclin C, Wijnholds J, Le Bivic A: PATJ connects and stabilizes apical and lateral components of tight junctions in human intestinal cells. J Cell Sci 2005, 118(Pt 17):4049–4057.

12. Fahey-Lozano N, La Marca JE, Portela M, Richardson HE: Drosophila Models of Cell Polarity and Cell Competition in Tumourigenesis. Adv Exp Med Biol 2019, 1167:37–64.

13. Mola-Caminal M, Carrera C, Soriano-Tarraga C, Giralt-Steinhauer E, Diaz-Navarro RM, Tur S, Jimenez C, Medina-Dols A, Cullell N, Torres-Aguila NP et al: PATJ Low Frequency Variants Are Associated With Worse Ischemic Stroke Functional Outcome. Circ Res 2019, 124(1):114–120.

14. Dashti HS, Daghlas I, Lane JM, Huang Y, Udler MS, Wang H, Ollila HM, Jones SE, Kim J, Wood AR et al: Genetic determinants of daytime napping and effects on cardiometabolic health. Nat Commun 2021, 12(1):900.

15. Wang H, Lane JM, Jones SE, Dashti HS, Ollila HM, Wood AR, van Hees VT, Brumpton B, Winsvold BS, Kantojarvi K et al: Genome-wide association analysis of self-reported daytime sleepiness identifies 42 loci that suggest biological subtypes. Nat Commun 2019, 10(1):3503.

16. Medina-Dols A, Canellas G, Capo T, Sole M, Mola-Caminal M, Cullell N, Jaume M, Nadal-Salas L, Llinas J, Gomez L et al: Role of PATJ in stroke prognosis by modulating endothelial to mesenchymal transition through the Hippo/Notch/PI3K axis. Cell Death Discov 2024, 10(1):85.

17. Ernkvist M, Persson NL, Audebert S, Lecine P, Sinha I, Liu M, Schlueter M, Horowitz A, Aase K, Weide T et al: The Amot/Patj/Syx signaling complex spatially controls RhoA GTPase activity in migrating endothelial cells. Blood 2009, 113(1):244–253.

18. Brenner S: The genetics of Caenorhabditis elegans. Genetics 1974, 77(1):71–94.

19. Castiglioni VG, Ramalho JJ, Kroll JR, Stucchi R, van Beuzekom H, Schmidt R, Altelaar M, Boxem M: Identification and characterization of Crumbs polarity complex proteins in Caenorhabditis elegans. J Biol Chem 2022, 298(4):101786.

20. Jiang WI, De Belly H, Wang B, Wong A, Kim M, Oh F, DeGeorge J, Huang X, Guang S, Weiner OD, Ma DK: Early-life stress triggers long-lasting organismal resilience and longevity via tetraspanin. Sci Adv 2024, 10(4):eadj3880.

21. Ritchie ME, Phipson B, Wu D, Hu Y, Law CW, Shi W, Smyth GK: limma powers differential expression analyses for RNA-sequencing and microarray studies. Nucleic Acids Research 2015, 43(7):e47–e47.

22. Wickham H: ggplot2: elegant graphics for data analysis, Second edition edn. Switzerland: Springer; 2016.

23. Siletti K, Hodge R, Mossi Albiach A, Lee KW, Ding S-L, Hu L, Lönnerberg P, Bakken T, Casper T, Clark M et al: Transcriptomic diversity of cell types across the adult human brain. Science 2023, 382(6667):eadd7046.

24. Yao Z, van Velthoven CTJ, Kunst M, Zhang M, McMillen D, Lee C, Jung W, Goldy J, Abdelhak A, Aitken M et al: A high-resolution transcriptomic and spatial atlas of cell types in the whole mouse brain. Nature 2023, 624(7991):317–332.

25. Zhu K, Bendl J, Rahman S, Vicari JM, Coleman C, Clarence T, Latouche O, Tsankova NM, Li A, Brennand KJ et al: Multi-omic profiling of the developing human cerebral cortex at the single-cell level. Science Advances 2023, 9(41):eadg3754.

26. Shin K, Wang Q, Margolis B: PATJ regulates directional migration of mammalian epithelial cells. EMBO Rep 2007, 8(2):158–164.

27. Lee YM: RUNX Family in Hypoxic Microenvironment and Angiogenesis in Cancers. Cells 2022, 11(19).

28. Ren R, Ding S, Ma K, Jiang Y, Wang Y, Chen J, Wang Y, Kou Y, Fan X, Zhu X et al: SUMOylation Fine-Tunes Endothelial HEY1 in the Regulation of Angiogenesis. Circ Res 2024, 134(2):203–222.

29. Liu S, Costa M: The role of NUPR1 in response to stress and cancer development. Toxicol Appl Pharmacol 2022, 454:116244.

30. Fang J, Luo S, Lu Z: HK2: Gatekeeping microglial activity by tuning glucose metabolism and mitochondrial functions. Mol Cell 2023, 83(6):829–831.

31. del Zoppo GJ, Hallenbeck JM: Advances in the vascular pathophysiology of ischemic stroke. Thromb Res 2000, 98(3):73–81.

32. Takata F, Nakagawa S, Matsumoto J, Dohgu S: Blood-Brain Barrier Dysfunction Amplifies the Development of Neuroinflammation: Understanding of Cellular Events in Brain Microvascular Endothelial Cells for Prevention and Treatment of BBB Dysfunction. Front Cell Neurosci 2021, 15:661838.

33. Abdullahi W, Tripathi D, Ronaldson PT: Blood-brain barrier dysfunction in ischemic stroke: targeting tight junctions and transporters for vascular protection. Am J Physiol Cell Physiol 2018, 315(3):C343–C356.

34. Fagan SC, Hess DC, Hohnadel EJ, Pollock DM, Ergul A: Targets for vascular protection after acute ischemic stroke. Stroke 2004, 35(9):2220–2225.

35. Williamson MR, Franzen RL, Fuertes CJA, Dunn AK, Drew MR, Jones TA: A Window of Vascular Plasticity Coupled to Behavioral Recovery after Stroke. J Neurosci 2020, 40(40):7651–7667.

36. Straight SW, Shin K, Fogg VC, Fan S, Liu CJ, Roh M, Margolis B: Loss of PALS1 expression leads to tight junction and polarity defects. Mol Biol Cell 2004, 15(4):1981–1990.

37. Hakanen J, Ruiz-Reig N, Tissir F: Linking Cell Polarity to Cortical Development and Malformations. Front Cell Neurosci 2019, 13:244.

38. Wang WJ, Lyu TJ, Li Z: Research Progress on PATJ and Underlying Mechanisms Associated with Functional Outcomes After Stroke. Neuropsychiatr Dis Treat 2021, 17:2811–2818.

39. Yu M, Jiang X, Cai W, Yang X, An W, Zhang M, Xu Y, Zhang B, Tang S: PATJ and MPDZ are required for trophectoderm lineage specification in early mouse embryos. Reproduction 2023, 166(2):175–185.

40. Lemmers C, Medina E, Delgrossi MH, Michel D, Arsanto JP, Le Bivic A: hINADl/PATJ, a homolog of discs lost, interacts with crumbs and localizes to tight junctions in human epithelial cells. J Biol Chem 2002, 277(28):25408–25415.

41. Moleirinho S, Hoxha S, Mandati V, Curtale G, Troutman S, Ehmer U, Kissil JL: Regulation of localization and function of the transcriptional co-activator YAP by angiomotin. Elife 2017, 6.

42. Boopathy GTK, Hong W: Role of Hippo Pathway-YAP/TAZ Signaling in Angiogenesis. Front Cell Dev Biol 2019, 7:49.

43. Wang X, Freire Valls A, Schermann G, Shen Y, Moya IM, Castro L, Urban S, Solecki GM, Winkler F, Riedemann L et al: YAP/TAZ Orchestrate VEGF Signaling during Developmental Angiogenesis. Dev Cell 2017, 42(5):462–478 e467.

44. Kim J, Kim YH, Kim J, Park DY, Bae H, Lee DH, Kim KH, Hong SP, Jang SP, Kubota Y et al: YAP/TAZ regulates sprouting angiogenesis and vascular barrier maturation. J Clin Invest 2017, 127(9):3441–3461.

45. Choi HJ, Zhang H, Park H, Choi KS, Lee HW, Agrawal V, Kim YM, Kwon YG: Yes-associated protein regulates endothelial cell contact-mediated expression of angiopoietin-2. Nat Commun 2015, 6:6943.

46. Sakabe M, Fan J, Odaka Y, Liu N, Hassan A, Duan X, Stump P, Byerly L, Donaldson M, Hao J et al: YAP/TAZ-CDC42 signaling regulates vascular tip cell migration. Proc Natl Acad Sci U S A 2017, 114(41):10918–10923.

47. Elaimy AL, Mercurio AM: Convergence of VEGF and YAP/TAZ signaling: Implications for angiogenesis and cancer biology. Sci Signal 2018, 11(552).

48. Hooglugt A, van der Stoel MM, Boon RA, Huveneers S: Endothelial YAP/TAZ Signaling in Angiogenesis and Tumor Vasculature. Front Oncol 2020, 10:612802.

49. Shen Y, Wang X, Liu Y, Singhal M, Gurkaslar C, Valls AF, Lei Y, Hu W, Schermann G, Adler H et al: STAT3-YAP/TAZ signaling in endothelial cells promotes tumor angiogenesis. Sci Signal 2021, 14(712):eabj8393.

50. Azad T, Ghahremani M, Yang X: The Role of YAP and TAZ in Angiogenesis and Vascular Mimicry. Cells 2019, 8(5).

51. Park JA, Kwon YG: Hippo-YAP/TAZ signaling in angiogenesis. BMB Rep 2018, 51(3):157–162.

52. Ong YT, Andrade J, Armbruster M, Shi C, Castro M, Costa ASH, Sugino T, Eelen G, Zimmermann B, Wilhelm K et al: A YAP/TAZ-TEAD signalling module links endothelial nutrient acquisition to angiogenic growth. Nat Metab 2022, 4(6):672–682.

53. Wigerius M, Quinn D, Fawcet JP: Emerging roles for angiomotin in the nervous system. Science Signaling 2020, 13(655):eabc0635.

54. Beck H, Plate KH: Angiogenesis after cerebral ischemia. Acta Neuropathol 2009, 117(5):481–496.

55. Hayashi T, Noshita N, Sugawara T, Chan PH: Temporal profile of angiogenesis and expression of related genes in the brain after ischemia. J Cereb Blood Flow Metab 2003, 23(2):166–180.

56. Font MA, Arboix A, Krupinski J: Angiogenesis, neurogenesis and neuroplasticity in ischemic stroke. Curr Cardiol Rev 2010, 6(3):238–244.

57. Trimm E, Red-Horse K: Vascular endothelial cell development and diversity. Nat Rev Cardiol 2023, 20(3):197–210.

58. Bonaguidi MA, Peng CY, McGuire T, Falciglia G, Gobeske KT, Czeisler C, Kessler JA: Noggin expands neural stem cells in the adult hippocampus. J Neurosci 2008, 28(37):9194–9204.

59. Ma X, Renda MJ, Wang L, Cheng EC, Niu C, Morris SW, Chi AS, Krause DS: Rbm15 modulates Notch-induced transcriptional activation and affects myeloid differentiation. Mol Cell Biol 2007, 27(8):3056–3064.

60. Bianchi ME, Mezzapelle R: The Chemokine Receptor CXCR4 in Cell Proliferation and Tissue Regeneration. Front Immunol 2020, 11:2109.

